# Modelling neuroanatomical variation due to age and sex during childhood and adolescence

**DOI:** 10.1101/126441

**Authors:** Gareth Ball, Chris Adamson, Richard Beare, Marc L. Seal, the Pediatric Imaging, Neurocognition and Genetics

## Abstract

Brain development is a dynamic process that follows a well-defined trajectory during childhood and adolescence with tissue-specific alterations that reflect complex and ongoing biological processes. Accurate identification and modelling of these anatomical processes *in vivo* with MRI may provide clinically useful imaging markers of individual variability in development. In this study, we build a model of age- and sex-related anatomical variation using multiple imaging metrics and manifold learning.

Using publicly-available data from two large, independent developmental cohorts (n=768, 862), we apply a multi-metric machine learning approach combining measures of tissue volume, cortical area and cortical thickness into a low-dimensional data representation.

We find that neuroanatomical variation due to age and sex can be captured by two orthogonal patterns of brain development and we use this model to simultaneously predict age with a mean error of 1.5-1.6 years and sex with an accuracy of 81%.

We present a framework for modelling anatomical development during childhood using manifold embedding. This model accurately predicts age and sex based on image-derived markers of cerebral morphology and generalises well to independent populations.

## Introduction

Brain development is a dynamic process that follows a well-defined trajectory during childhood and adolescence. During this formative period the brain undergoes profound change: increases in brain size, most rapid after birth, continue into late adolescence (Dekaban and Sadowsky, 1978); myelination processes that begin *in utero* continue to progress through to the second decade of life (Yakovlev and Lecours, 1967), and synaptic pruning leads to significant reductions in synaptic density during early adolescence (Huttenlocher, 1979).

Magnetic resonance imaging (MRI) provides the opportunity to study brain development and track developmental processes in vivo. Analyses of structural MRI have found that grey and white matter volumes follow different trajectories during adolescence. Cortical grey matter volume is greatest in childhood, then gradually decreases during adolescence (Mills et al., 2016; Tamnes et al., 2013), whereas white matter volume increases throughout childhood and adolescence (Aubert-Broche et al., 2013; Lebel and Beaulieu, 2011). In contrast, measures of cortical thickness show progressive thinning from mid-childhood through to the early 20s (Raznahan et al., 2011) whereas cortical surface area appears to follow a cubic trajectory, peaking in late childhood before declining through adolescence (Wierenga et al., 2014). These observations were recently confirmed across four independent developmental samples, where Tamnes et al. observed consistent developmental trajectories characterised by a decreasing cortical thickness with increasing age and childhood increases in surface area followed by subtle decreases during adolescence (Tamnes et al., 2017). Beyond these global measures, significant regional differences in development have also been reported (Ducharme et al., 2016; Gogtay et al., 2004; Vijayakumar et al., 2016). Fjell et al. found that areal expansion in several cortical regions was greater than that of the average increase across the whole cortex; these regions included the anterior cingulate cortex, frontal cortex and insula (Fjell et al., 2015). Similarly, regional measures of cortical volume show a differential effect of age, with changes in some regions (e.g.: medial parietal cortex) most apparent at younger ages, compared to others where the rate of change was greatest later in development (e.g.: the anterior temporal lobe) (Tamnes et al., 2013). Sex also likely plays a role in cerebral development. Sexual dimorphism has been observed in developmental studies of cortical thickness (Sowell et al., 2007) and sex-by-age interactions in area and folding have been reported in frontal and temporal cortex in adolescence (Koolschijn and Crone, 2013; Mutlu et al., 2013). Conversely, other studies suggest that cortical volumes follow a similar trajectory, but reach a ‘peak’ later in males than in females, the timing of which may coincide with pubertal onset (Giedd et al., 1999; Lenroot et al., 2007), although the accurate definition of developmental peaks is difficult and may be at risk to potential model or sample biases (Fjell et al., 2010; Mills and Tamnes, 2014). In addition, male brains are larger than females throughout development (Paus et al., 2017), and regional, volumetric estimates of sex differences may also be diminished once corrected for global differences in scale (Mills et al., 2016).

Taken together, these studies present a consensus view of typical cerebral development. However, longitudinal studies of healthy populations have shown that individual development can deviate significantly from these canonical trajectories (Mills and Tamnes, 2014), highlighting the need to create models of typical growth and development of the brain that can be applied on an individual level. Establishing developmental trajectories for typical cerebral development is vital to our understanding of maturational brain change and may provide a more accurate understanding of relationships between brain maturation and behavioural phenotypes during development.

Recently, studies have shown that it is possible to use MRI to accurately predict an individual’s age (Brown et al., 2012; Dosenbach et al., 2010; Erus et al., 2015; Franke et al., 2010; Khundrakpam et al., 2015). These methods involve the use of modern machine-learning techniques to extract informative morphological features to act as markers of cerebral maturation in a predictive model. Using such models, the difference between chronological age and age predicted from imaging has been framed as an index of accelerated or delayed development or aging, depending on the direction of the discrepancy (Cole et al., 2015; Franke et al., 2012; Löwe et al., 2016). For example, Franke et al. used a nonlinear, multivariate kernel regression model to predict age in a cohort of children and adolescents based on maps of grey matter volume (Franke et al., 2012). They were able to predict age accurately across the full age-range. Moreover, predicted brain age was found to be significantly lower in a small clinical population of preterm-born adolescents, suggesting a developmental delay in brain maturation (Franke et al., 2012). Recently, Erus et al. reported that a significant increase in predicted brain age compared to chronological age in childhood (predicted by a multi-modal MRI assessment) was indicative of precocious cognitive development (and *vice versa*), suggesting that complex cognitive phenotypes could be captured as variation along a single dimension of brain development (Erus et al., 2015).

In this study, we aim to build upon these studies by constructing a model of typical brain development during childhood and adolescence using manifold learning, exploiting a rich data source of MRI acquired in a large-scale developmental cohort (Jernigan et al., 2016). Manifold learning refers to a suite of dimension reduction techniques based on the intuition that high-dimensional datasets (such as MRI) reside on an embedded low-dimensional, and possibly non-linear, manifold or subspace. As such it represents a natural setting for this problem. The aim is to learn a mapping between the high- and low-dimensional data representations while preserving certain statistical properties (e.g.: variance) of the original data. This form of dimension reduction allows for an easier and more intuitive interpretation of important model features while retaining the underlying nonlinear relationships present between individual datapoints in the original dataset.

Here, we introduce the use of neighbourhood-preserving embedding (NPE) as a method to isolate structured patterns of covariance within populations, simultaneously modelling typical neuroanatomical variation due to age and sex during development as a low-dimensional process. Using a multi-metric approach, we combine measures of tissue volume, cortical area and cortical thickness to build a model that predicts age and sex with high accuracy and generalises well to other developmental populations. We also test the hypothesis that deviation from this model is associated with cognitive performance in two large population-based cohorts.

## Methods

### Imaging data

To model typical neurodevelopment, 3 Tesla, T1-weighted MRI data were obtained from the PING Study (Jernigan et al., 2016). The PING cohort comprises a large, typically-developing paediatric population with participants from several US sites included across a wide age and socioeconomic range. Exclusion criteria included: a) neurological disorders; b) history of head trauma; c) preterm birth (less than 36 weeks); d) diagnosis of an autism spectrum disorder, bipolar disorder, schizophrenia, or mental retardation; e) pregnancy; and f) daily illicit drug use by the mother for more than one trimester (Jernigan et al., 2016). Similar proportions of males and females participated across the entire age range.

The PING cohort included 1493 participants aged 3 to 21 years, of whom 1249 also had neuroimaging data. Of these, n=773 were available to download from NITRC (https://www.nitrc.org). T1 images were acquired using standardized high-resolution 3D RF-spoiled gradient echo sequence with prospective motion correction (PROMO) at 9 different sites, with pulse sequences optimized for equivalence in contrast properties across scanner manufacturers (GE, Siemens, and Phillips) and models (for details, see Jernigan et al., 2016). Written parental informed consent was obtained for all PING subjects below the age of 18 and directly from all participants aged 18 years or older.

In addition to imaging data, participants undertook comprehensive behavioural and cognitive assessments (NIH Toolbox Cognition Battery, NTCB; Akshoomoff et al., 2014) and provided a saliva sample for genome-wide genotyping. The NTCB comprises seven tests (Flanker, Picture Sequence, List Sorting, Picture Vocabulary, Reading, Dimensional Change Card Sorting, Pattern Comparison) that measure abilities across six major cognitive domains, including cognitive flexibility, inhibitory control, and working memory (Akshoomoff et al., 2014).

For model validation, a comparative neurodevelopmental population was obtained from the ABIDE and ABIDEII datasets (Di Martino et al., 2017, 2014). These datasets represent a consortium effort to aggregate MRI datasets from individuals with autism spectrum disorder and age-matched typically-developing controls. Contributions per site ranged from 13 to 105 typically-developing participants per site chosen as matched controls for the ASD population at each site. For both studies, 3 Tesla, T1-weighted MRI were acquired from 17 sites; images and acquisition details are available at http://fcon_1000.projects.nitrc.org/indi/abide. All participating sites received local Institutional Review Board approval for acquisition of the contributed data. In addition to imaging data, phenotypic information including age, sex, IQ and diagnostic information were recorded and made available (Di Martino et al., 2017, 2014)

### Image processing

Quality control assessment for the PING data is detailed in Jernigan et al. (2016). In brief, images were inspected for excessive distortion, operator compliance, or scanner malfunction. Specifically, T1-weighted images were examined slice-by-slice for evidence of motion artefacts or ghosting and rated as acceptable, or recommended for re-scanning. After additional, on-site, visual quality control assessment, we removed a further 5 participants, resulting in a final sample of n=768 (mean age=12.3y; range: 3.2–21.0y; 404 male). Site-specific demographic data are shown in Table S1.

The ABIDE cohort included imaging data from 1112 participants. After initial visual quality assessment, 18 were removed due to motion artefacts and 10 due to poor image contrast. A further 28 were removed due to prior pre-processing (n=1), or high similarity to other images within ABIDE, or ABIDE-II (i.e.: repeat scans). Remaining T1 images from typically-developing participants aged 21y and under from 17 sites were included in the final sample of n=424 (346 male; mean age=13.69y).

In total, n=1044 datasets were available to download from ABIDE-II. Of these, 28 were excluded due to visible image artefacts, and a further 13 failed cortical reconstruction. Two images appeared to have corresponding repeat scans and were removed. As above, we removed those aged over 21, and those with an ASD diagnosis, resulting in a final sample of n=439 (297 male; mean age=11.50y) participants from 15 sites. Site-specific demographic data for ABIDE and ABIDE-II are shown in Table S2.

For all subjects, vertex-wise maps of cortical thickness and cortical area (estimated on the white matter surface) were constructed from T1 MRI with FreeSurfer 5.3 (http://surfer.nmr.mgh.harvard.edu). Briefly, this process includes removal of non-brain tissue, transformation to Talairach space, intensity normalisation, tissue segmentation and tessellation of the grey matter/white matter boundary followed by automated topology correction. Cortical geometry was matched across individual surfaces using spherical registration (Dale et al., 1999; Fischl et al., 2002, 1999; Fischl and Dale, 2000). Any images that failed initial surface reconstruction, or returned surfaces with topological errors, were manually fixed using white matter mask editing and re-submitted to Freesurfer until all datasets passed inspection. In total, 13 ABIDE and 16 ABIDE-II images required editing to complete surface reconstruction.

In addition, whole-brain tissue volume maps were estimated using deformation-based morphometry (Ashburner et al., 1998; Rueckert et al., 2003). Each participant’s T1 image was intensity normalised, corrected for bias field inhomogeneities and aligned to MNI 152 space using diffeomorphic nonlinear registration (ANTs; Avants et al., 2008; Tustison et al., 2010). Transformed images were visually inspected to ensure alignment to the template images and voxel-wise maps of volume change induced by the nonlinear deformation were characterised by the determinant of the Jacobian operator, referred to here as the Jacobian map. Each map was log-transformed so that values greater than 0 represent local areal expansion in the subject relative to the target and values less than 0 represent areal contraction.

Prior to analysis, both tissue volume maps and cortical thickness and area maps were smoothed with a Guassian kernel of 10 FWHM.

### Manifold learning

Neighbourhood preserving embedding (NPE) is a linear approximation to locally linear embedding (LLE; Roweis and Saul, 2000) that seeks to find a linear transformation, *P*, to map a high-dimensional *n* × *D* dataset *X* = {*X*_1_, *X*_2_,…, *X_n_*} into a low-dimensional *n* × *d* subspace *Y* = [*Y*_1_, *Y*_2_,…, *Y_n_*} where *d* ≪ *D* and *Y* = *P^T^X* (He et al., 2005). Like LLE, and in contrast to other linear subspace methods (e.g.: PCA), NPE aims to preserve the local neighbourhood structure of the data. That is, communities, patterns, or between-group differences that exist among datasets in the high-dimensional setting should be preserved within the low-dimensional subspace.

The process is illustrated in Fig 1A. For a given data point *X_i_*, an adjacency matrix is first constructed, placing an edge between *X_i_* and *X_j_* only if *X_j_* belongs to the set of *k* nearest neighbours of *X_i_*. Following this, a set of weights, *W,* is calculated that approximately reconstruct *X_i_* from its neighbours and a linear projection, *P,* sought to optimally preserve this neighbourhood structure in the low-dimensional space, *Y* (He et al., 2005). One of the major benefits the NPE confers over LLE is that the solution generalises to new datapoints, allowing unseen data to be projected onto the manifold without re-calculating the embedding.

Local neighbourhoods are typically defined based on the Euclidean distance between samples in the high-dimensional image space, however it is possible to introduce other constraints in order to conduct NPE in a supervised setting (Fig. 1B; He et al., 2005; Zeng and Luo, 2007). For example, restricting local neighbourhoods to only include subjects from the same class to enhance group separation, or weighting similarities based on some other subject-specific attributes of interest (e.g.: age).

**Figure 1:**
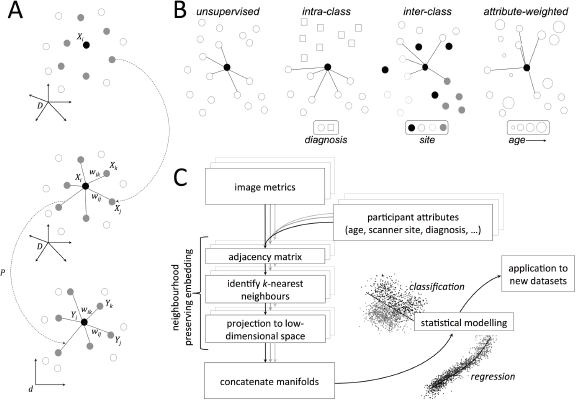
Neighbourhood preserving embedding and image analysis pipeline. A. For a given datapoint, *X_i_*, nearest neighbours are selected and weights assigned that can be used to approximately reconstruct *X_t_*. A linear transformation, *P*, is then sought to project the data into a low-dimensional space while preserving the neighbourhood structure. B. Possible supervision strategies for neighbourhood construction. In an unsupervised setting, neighbours are selected based on image similarity alone; alternatively, neighbours can be selected from within- or between-classes in order to maximise/minimise group differences in the manifold structure. Similarly, neighbours can be selected based on the weighted combination of image similarity and that of another subject-specific attribute (e.g.: age). C. Analysis pipeline for NPE analysis. For each image metric, NPE is used for subspace projection, before the embedded data are combined and passed on for statistical modelling.

The analysis pipeline used in this study is shown in Fig. 1C. For each image metric (tissue volume, cortical thickness, cortical area), data were first mean-centred and projected to an orthogonal subspace via singular value decomposition (SVD), while retaining 95% of variance. NPE was then performed using ***k*** = 10 neighbours, projecting data to ***d*** = 3 dimensions. In order to maximally preserve age- and sex-related variation in the embedded data, we incorporated participant attributes into the construction of the adjacency matrices. Nearest neighbours were selected based on the product of two adjacency matrices, *A* and *a*, where the *(i, j)^th^* element of each matrix represents the (normalised) similarity between images, *A,* and age, *a*, of subjects *i* and *j*, respectively and *A_i,j_* = 0, if *S_i_* ≠ *S_j_*, where *S* indicates the sex of the participant. In order to reduce potential bias in image similarities due to site effects, we also introduced an additional constraint: *A_i,j_* = 0, if *S_i_* = *S_j_*, where 5 indicates the site/ scanner of image acquisition, although this had little effect on the final embedding.

This resulted in three sets of coordinates, *Y_v_*, *Y_t_*, and *Y_a_*, representing the low-dimensional embedding of tissue volume, *V*, cortical thickness, *t*, and area, *a,* data. This produced a final low-dimensional representation of the combined, multi-metric image data for statistical analysis.

Example code for performing supervised NPE is made available at http://developmentalimagingmcri.github.io.

### Statistical analysis

Internal validity of the model was assessed using 10-fold cross-validation. We used 90% of the PING participants as a training set, calculating the manifold embedding coordinates for each imaging modality and combining them into a single representation. To predict age, we used a Gaussian Process Regression (GPR) model with the combined manifold coordinates as features and age as a dependent variable (Cole et al., 2015). To predict sex, the combined coordinate set was sent to a linear discriminant classifier. Image data from the remaining 10%, the test set, were then projected onto the joint manifold and the fitted models used to predict age and sex. Mean absolute error in age estimation (MAE) and correlation between true age and predicted age are reported, alongside classification accuracies for sex. This process was repeated for each fold, reconstructing the manifold each time, such that all PING participants were part of the test set exactly once.

External validity was assessed by projecting the combined ABIDE and ABIDE-II datasets onto a manifold constructed from the full PING dataset and predicting age and sex using models trained on the PING data.

To determine if errors in image-derived age estimation correlated with cognitive performance, we performed a further set of analyses using available cognitive data. Of the PING dataset, n=617 had complete records for NTCB score, family socioeconomic status (household income and parental education), and genetic ancestry (GAF; Jernigan et al., 2016; Libiger and Schork, 2012). NTCB scores were corrected for age, sex, socioeconomic status and GAF (Akshoomoff et al., 2014) and linear regression used to determine associations between age estimation error (based on cross-validated age predictions) and corrected cognitive scores.

For validation, in the ABIDE and ABIDE-II datasets, full scale IQ (FSIQ) was available for n=802 typically-developing participants (out of 863 combined across both studies). Using the same 10-fold cross-validation procedure outlined above, we derived age predictions for each participant and performed linear regression between age estimation error and FSIQ.

All statistical analysis was performed in Matlab R2105b (Natick, MA).

## Results

Figure 2 shows manifold structure visualised for tissue volume (Fig 2A), cortical area (Fig 2B) and cortical thickness (Fig 2C) calculated in the PING cohort. For each metric, an orthogonal rotation was applied to each manifold to maximise correlation with age and sex along the first and second axes respectively. The images show the (standardised) weight of the embedding vectors (model coefficients) used to project new data to the rotated manifold, thus important features are represented by a larger weight, and increases in volume, thickness or area in regions with high positive weight will result in a positive increase along the respective embedding axis and *vice versa.*

**Figure 2.**
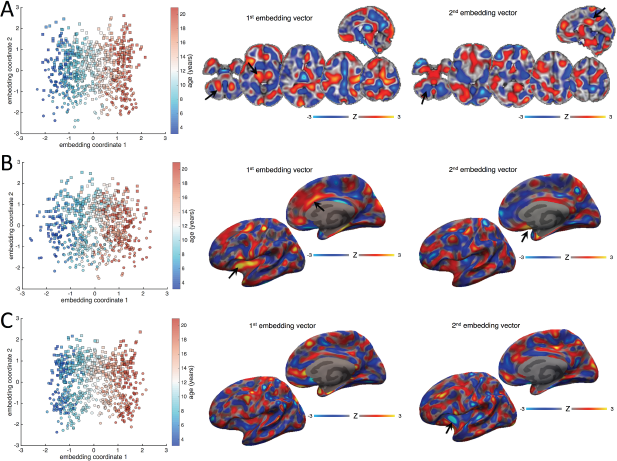
Manifold structure for tissue volume, cortical area and cortical thickness. Manifold structure is visualised for tissue volume (A), cortical area (B) and cortical thickness (C). For each image metric, the first two embedding coordinates are plotted against each other. Each point represents a subject; the colourbar indicates age and markers denote sex (square: male; circle: female). Images show the embedding vectors for the first and second coordinates, i.e.: the model coefficients in each voxel required to transform data into the embedded subspace. Maps are Z-scored for comparison (colourbar).

For all three modalities, age increases along the first embedding coordinate, indicated by the gradation of colour along the first axis of the scatter plots Increasing age (a positive embedding coordinate) is associated with a neuroanatomical pattern represented by relatively increased tissue volume in the cerebellum, brain stem and thalamus (black arrows; Fig 2A), and ascending white matter tracts subjacent to the primary motor cortex (positive image weights), with relative decreases in medial frontal and parietal cortices (negative image weights, Fig 2A). In the cortex, age is associated with increasing surface area in the insula and anterior cingulate cortex (black arrows, Fig 2B).

Separation between sexes is shown along the second dimension (squares and circles, Fig 2). This is associated with a distributed, discriminant pattern of tissue volume alterations with increases in medial posterior cingulate (black arrow; Fig 2A) and primary visual cortex, and brainstem, and relative decreases in the basal ganglia, frontal pole, and cerebellum (negative weights, black arrow) associated with male sex. Separation along the second dimension was also associated with regions of increased surface area in medial temporal and orbitofrontal cortices (black arrow, Fig 2B), and decreased cortical thickness in the anterior insula (arrow, Fig 2C).

Figure 3A shows the joint manifold structure visualised after concatenating all three image metrics and performing a final dimension reduction on the concatenated coordinate data, Y_c_ = (Y_v_, Y_t_, Y_a_) using PCA (Aljabar et al., 2011). This demonstrates how two orthogonal patterns of anatomical variation associated with age and sex during development can be captured along two dimensions in this population.

**Figure 3.**
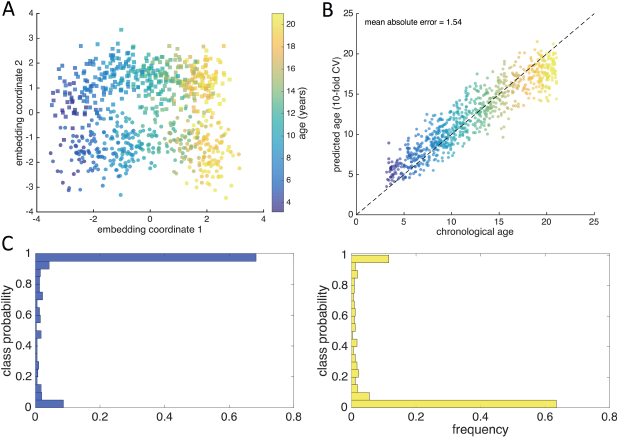
Age and sex prediction with manifold embedding. A. The first two coordinates of the joint manifold are shown, each point represents a subject; the colourbar indicates age and markers denote sex (square: male; circle: female). B. Using 10-fold cross-validation, age and sex were predicted using on the first two embedding coordinates. Gaussian Process regression was used to predict age, shown plotted against chronological age (colourbar shown as in A). C. Predicted class probabilities are shown for males (blue histogram) and females (yellow).

Using 10-fold cross-validation, we predicted age in the PING cohort with an MAE of 1.54 years (correlation between chronological and predicted age=0.926; Fig 3B). Using a linear discriminant classifier, we predicted sex with an accuracy of 80.9% (Fig 3C). We repeated this analysis, accounting for differences in global scale by correcting voxel-(vertex-)wise measures of tissue volume, cortical surface area and thickness metrics for intracranial volume, total surface area and mean cortical thickness, respectively. Manifold structure for each metric is shown in Fig S1 after global correction, inspection of the embedding vectors revealed similar patterns to those shown in Fig 2. After correcting for global scale, we achieved a cross-validated MAE of 1.79 years, and an accuracy of 71.4% in the PING cohort.

To determine if model accuracy varied with age, we partitioned our data into 10 approximately equal-sized bins and calculated MAE and classification accuracy in each (Table 1). Age prediction error ranged from a minimum of 1.23y at around 9 years of age, to a maximum of 2.55y in the oldest participants (mean age=20.3y). In contrast, classification accuracy ranged from 69% to 87%, with discrimination lowest in the youngest participants (mean age = 4.5y) and highest at around 16 years. After correction for global scale, the minimum MAE was 1.50 (mean age 11.7y; maximum MAE=2.9, mean age 22.3y), and the lowest and highest classification accuracies were 65.3% in the youngest group, and 80.3% at 16 years.

**Table 1.** Age and sex prediction accuracy at different ages

| Bin | n | mean age | MAE | male (%) | accuracy |
| --- | --- | --- | --- | --- | --- |
| 1 | 75 | 4.48 | 1.69 | 38 (50.1) | 69.33 |
| 2 | 79 | 7.31 | 1.32 | 44 (55.7) | 79.75 |
| 3 | 71 | 8.96 | 1.33 | 38 (53.5) | 78.87 |
| 4 | 79 | 9.63 | 1.23 | 35 (44.3) | 82.28 |
| 5 | 77 | 11.72 | 1.40 | 43 (55.8) | 81.82 |
| 6 | 80 | 12.89 | 1.46 | 52 (65.0) | 82.50 |
| 7 | 77 | 14.52 | 1.29 | 47 (61.0) | 81.82 |
| 8 | 76 | 16.15 | 1.46 | 36 (47.4) | 86.84 |
| 9 | 74 | 17.63 | 1.62 | 34 (46.0) | 81.08 |
| 10 | 80 | 20.29 | 2.55 | 38 (47.5) | 83.75 |

To demonstrate external validity of this approach, we repeated this analysis using a model constructed from the full PING dataset to predict age and sex in the ABIDE and ABIDE-II cohorts (Fig 4). We predicted age in ABIDE with an MAE of 1.65 years (correlation: 0.825), and sex with an accuracy of 80.0% (1.87y and 71.2% after global correction). We achieved similar results in ABIDE-II (MAE=1.59, correlation: 0.817; accuracy=80.2; 1.80y and 70.6% after global correction).

**Figure 4.**
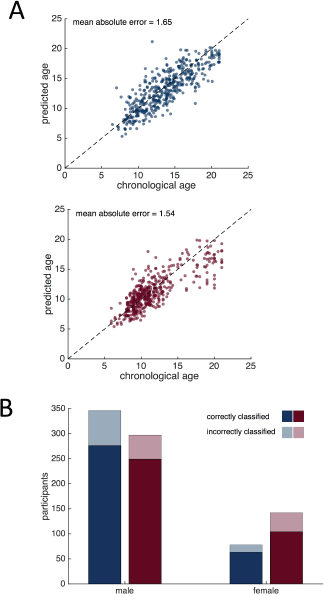
Age and sex prediction in typically-developing ABIDE and ABIDE-II participants. A. Scatterplots show ABIDE (blue) and ABIDE-II (red) data projected onto the manifold constructed from the full PING dataset. Line indicates identity (i.e.: predicted = chronological age). B. Correctly and incorrectly classified male and female participants in both groups are shown.

Prediction accuracies in the PING cohort were robust to altering the number of neighbours, *k*, used in manifold construction (*k* =5: classification accuracy=80.9%, MAE=1.51; *k* =20: accuracy =79.7; MAE=1.53), and the number of manifold dimensions, *d* (*d*=5: accuracy=80.1%, MAE=1.59; *d* =10: accuracy=81.1%, MAE=1.59). Our model was robust to site variation in the PING cohort: performing NPE without the additional site constraint did not affect the prediction accuracies (MAE=1.49y, accuracy=80.0%) and there was no significant association between acquisition site and absolute age estimation error (ANOVA: F_6,761_ = 1.95, p=0.07; Fig S2). A main effect of site was evident in both ABIDE and ABIDE-II age prediction error (F16,407 = 1.86, p=0.02; F_14,424_ = 16.56, p<0.001; Fig S2), although this was likely exacerbated by an uneven age distributions between sites in this cohort (main effect of site on age: ABIDE F_16,407_=14.36, p<0.001; ABIDE-II F_14, 424_=68.4, p<0.001). Finally, we found that using NPE for subspace projection outperformed PCA (MAE = 1.61y, accuracy=71.2%; with global correction, MAE=1.91y, accuracy=54.4%).

To assess the individual contribution of each set of image metrics, we also performed the analysis using only tissue volume, cortical thickness or cortical area data. We found that the joint model combining all three metrics outperformed single metric models for both age and sex prediction (tissue volume only: MAE=1.69, accuracy=71.9%; area: MAE=2.77, accuracy=72.7%; thickness: MAE=2.07, accuracy=71.5%). Fig S2 shows MAE and accuracies for each tissue metric with and without correction for global scale.

### Associations with cognition

In order to determine if deviations from the average developmental trajectory of the brain coincided with cognitive performance we compared predicted age errors (the difference between age estimated from MRI using the above model and true, chronological age) with cognitive scores in PING and ABIDE.

In the PING cohort, no significant associations were found between NTCB scores (corrected for age, sex, socioeconomic status and genetic ancestry) and predicted age error after correcting for multiple comparisons (Table 2).

**Table 2:** Associations between predicted age error and cognitive score in PING

| NTCB score | Linear regression |  |  |
| --- | --- | --- | --- |
|  | R<sup>2</sup> | F<sub>1,615</sub> | p |
| Flanker | 0.000 | 0.018 | 0.892 |
| Attention | 0.000 | 0.068 | 0.794 |
| Picture Sequence Memory | 0.007 | 4.292 | 0.039 |
| List Sorting | 0.001 | 0.469 | 0.494 |
| Picture Vocabulary | 0.003 | 2.118 | 0.146 |
| Reading | 0.000 | 0.072 | 0.789 |
| Dimensional Change Card Sorting | 0.001 | 0.315 | 0.575 |
| Pattern Comparison | 0.007 | 4.247 | 0.040 |

We repeated this analysis using predicted age estimates form 10-fold cross-validation in typically-developing participants from the ABIDE and ABIDE-II cohorts (n=802 with cognitive scores). Using full scale IQ as a measure of cognitive performance, we found no strong evidence for an association with predicted age error (F_1_,_800_=0.649, p=0.421). This remained the case when after correcting FSIQ for effects of site and sex (F_1,800_=0.103, p=0.748).

## Discussion

The brain follows a well-defined developmental trajectory during childhood, with tissue-specific alterations that reflect complex and ongoing biological processes including myelination and synaptic pruning. Accurate identification and modelling of these anatomical processes *in vivo* with MRI may provide clinically useful imaging markers of individual variability in development. In this study, we use manifold learning to generate a parsimonious description of typical brain development during childhood and adolescence. By combining measures of tissue volume, cortical thickness and cortical area, we show how patterns of anatomical variation can be used to accurately predict age and sex between the ages of 3 and 21 years. We show that this model is not strongly affected by site-to-site variation in image acquisition and yields accurate predictions across different study populations. In contrast to previous reports, however, we do not find strong evidence that deviation from a predicted trajectory corresponds to adverse functional or behavioural outcome in healthy individuals.

We demonstrate that biological age can be predicted from MRI in developmental populations with a mean error of around 1.6 years. This is in line with previous reports in this population. Using a set of 231 pre-selected, image-based features from T1, T2 and diffusion-weighted MRI, Brown et al. developed a nonlinear model of cerebral maturation to predict age in the PING cohort, achieving an MAE of 1.03 years (Brown et al., 2012). Using just T1-weighted image features, Brown et al. reported an MAE of 1.71y, comparable to the MAE reported in this study. Similarly, they also found that model accuracy decreased slightly with increasing age, suggesting that image-based prediction is most accurate during the periods when the rate of anatomical change is greatest (Brown et al., 2012; Dekaban and Sadowsky, 1978). Indeed, in both PING and the ABIDE cohorts, we observe a trend towards a model-based underestimation of age in the older participants. This error can be seen in other MRI-based age estimation methods (Han et al., 2014; Khundrakpam et al., 2015) and may indicate the increasing difficulty of discrimination between individuals in early adulthood compared to childhood and adolescence.

Using a similar approach, Franke et al. reported an MAE of 1.1 years in a developmental cohort aged 5 to 18 (Franke et al., 2012). Using nonlinear mapping functions (i.e.: kernels) in machine learning allows for the use of linear methods to discover highly nonlinear boundaries or patterns in the original data by creating an implicit feature space (Hofmann et al., 2008). While flexible, a limitation of these methods is the inability to identify important features in the space of the original dataset. An alternative approach may be to use regularised linear regression across all cortical regions, although this method depends upon an initial cortical parcellation (Khundrakpam et al., 2015). By calculating a linear mapping between the original, high-dimensional data and the low-dimensional embedded manifold, NPE produces a set of basis vectors that — through linear combination – can approximately reconstruct the original dataset, capture nonlinear relationships and provide interpretable voxel-(vertex-)wise maps of feature importance (He et al., 2005).

We show examples of these vectors in Figure 2 for each image metric. The images represent distributed patterns of neuroanatomical variation that highlight regional importance within the model. As reported previously, increasing age is reflected a lower-to-higher-order trajectory characterised by reduced tissue volume in frontal and parietal regions, coupled with increased white matter tissue and brain stem volume (Aubert-Broche et al., 2013; Sowell et al., 2004; Xie et al., 2012). Increasing age was also associated with increased cortical surface area most prominent in the insula and cingulate cortex. This agrees with previous reports of high rates of cortical surface area expansion during childhood in regions associated with higher-order intellectual function (Amlien et al., 2016; Fjell et al., 2015).

Sexual dimorphism during development is a contentious issue. Developmental trajectories for cortical grey and white matter appear similar between sexes (Aubert-Broche et al., 2013; Mills et al., 2016) with perceived sex differences often assigned to variation in physical size (Dekaban and Sadowsky, 1978; Giedd et al., 2012). In a longitudinal study of 387 subjects aged 3 to 27, Lenroot et al. reported increased frontal grey matter volume in females and increased occipital white matter in males, after accounting for brain size (Lenroot et al., 2007). Conversely, Sowell et al. reported thicker parietal and posterior temporal cortex in females, independent of age (Sowell et al., 2007). These discrepant findings may reflect the different timing of puberty or the differential effects of testosterone on brain development males and females (Bramen et al., 2012).

Here, we find that male sex is predicted by a pattern of neuroanatomical variation including increased brain tissue volume in the posterior cingulate and occipital lobe, volumetric decreases in the basal ganglia and insula, alongside a pattern of reduced cortical surface area in medial frontal regions and decreased cortical thickness in the insula. We highlight that the manifold embedding coordinates defined by NPE are orthogonal by construction; as such, the patterns shown in Figure 2 reflect anatomical variation independently associated with age and sex during development. After correcting for global differences in intracranial volume, total surface area and mean cortical thickness, we were still able to achieve relatively accurate predictions of age and sex across development, although both were maximised with the inclusion of global scaling information. Importantly, our study confirms recent reports that multivariate analyses that consider whole-brain patterns of variation in brain morphometry can reliably and accurately discriminate sex, even if large within-class, or between region, variability exists (Chekroud et al., 2016; Rosenblatt, 2016). We also show that this dimorphic pattern is evident even at very young ages, achieving a classification accuracy of around 69% (65% after global correction) in the youngest participants (aged 3-5y). It is important to consider that this finding does not imply that all females have e.g.: a smaller posterior cingulate than all males, or even that the cingulate is on average smaller in females across populations (see Fig S3 for an example of this). We suggest that this pattern of variation is one of a number (including the pattern of age-related variation described above) that exist concurrently within a population. An individual’s anatomical phenotype can then be viewed as arising from the weighted expression of these patterns, with the weight dependent on e.g.: age, sex, environmental or genetic factors. As such, tissue volume or cortical thickness, measured at a single point reflects the combination of multiple, distributed patterns of variation and, as such, may vary greatly over a population and not necessarily in line with a given covariate of interest (e.g.: sex).

Surprisingly, and in contrast to previous reports, we do not find strong evidence that a divergent image-based age prediction is a good marker of cognitive function. Using a multi-modal approach, that also included measures derived from diffusion MRI, the discrepancy between chronological age and an image-based estimate of ‘brain age’ has been proposed as a possible biomarker of cognitive development (Erus et al., 2015). This follows on from studies in older populations, where a more advanced ‘brain age’ was used as a marker of aging and found to correlate with brain injury (Cole et al., 2015) and dementia (Gaser et al., 2013). In this study, we tested this hypothesis in two large, independent, developmental cohorts. Using cross-validated age predictions, we did not find any statistically significant associations between measures of cognitive function (NCTB scores in PING; FSIQ in ABIDE) and age prediction error in typically-developing individuals. In the PING cohort, Akshoomoff’ et al., found that age, sex, socioeconomic status and genetic ancestry explained between 57% and 73% of variance in each of the NCTB scores (Akshoomoff et al., 2014). After correcting for these factors, we found that brain age estimation error explained at most 0.7% of the remaining variance (in memory and pattern comparison tests). In the ABIDE cohort, we found no associations between FSIQ and age estimation error. This suggests that model error in age prediction does not reflect the impact of an underlying latent variable associated with cognition. This discrepancy is likely due to differences in model construction between methods. Here, we use NPE to extract a single dimension of anatomical variation that aims to maximally preserve specific age-related structure in the data. As such, the reported pattern may lie orthogonal to neuroanatomical correlates of (non age-related) cognitive performance and as mentioned above, such patterns, while coexistent within the population, may not vary as a function of each other. Other methods that predict age based on the appearance of the brain as a whole, might better reflect the conflation of cognitive and age-related ‘components’ during development, such that model error captures variation in anatomy aligned with cognition (Erus et al., 2015).

Our findings suggest that the anatomical maturation of the brain during childhood and adolescence can be accurately modeled within a low-dimensional subspace. That is, variation along two axes is sufficient to capture individual variations due to age and sex within a population with relatively high accuracy. In contrast, functional or cognitive development is not well represented by variation along these axes. This suggests that additional, orthogonal dimensions of development are required to more accurately model individual trajectories. Alternatively, this approach may benefit from incorporating information from additional imaging modalities (e.g.: functional MRI) in order to more fully capture phenotypic variation associated with cognitive development (Erus et al., 2015; Liem et al., 2017). Subspace projection methods, such as NPE, bring focus to reliable and robust patterns that can predict phenotypic characteristics based on the brain’s shape and appearance. In a clinical setting, this framework could be extended to explore anatomical patterns underlying developmental or neuropsychiatric disorder, or stratifying clinical populations by locating individuals within clusters based on the expression of different neuroanatomical imaging components. Indeed, combining functional, diffusion and possibly genetic information into a larger manifold framework and considering similarities over multiple modalities to model local neighbourhoods and communities within large datasets could provide a more complete model of individual variation during this time period. In addition, the projection of longitudinal data onto the manifold could enable individuals to be tracked over time, an important consideration for developmental studies (Mills and Tamnes, 2014).

In summary, we present a framework for modelling anatomical development during childhood. This model accurately predicts age and sex based on image-derived markers of cerebral morphology and generalises well to independent populations.

## Acknowledgements

This research was conducted within the Developmental Imaging research group, Murdoch Childrens Research Institute and the Children’s MRI Centre, Royal Children's Hospital, Melbourne, Victoria. It was supported by the Murdoch Childrens Research Institute, the Royal Children’s Hospital, Department of Paediatrics, The University of Melbourne and the Victorian Government's Operational Infrastructure Support Program. The project was generously supported by RCH1000, a unique arm of The Royal Children’s Hospital Foundation devoted to raising funds for research at The Royal Children’s Hospital.

Data collection and sharing for this project was funded by the Pediatric Imaging, Neurocognition and Genetics Study (PING) (NIH: RC2DA029475). PING is funded by the National Institute on Drug Abuse and the Eunice Kennedy Shriver National Institute of Child Health & Human Development. PING data are disseminated by the PING Coordinating Center at the Center for Human Development, University of California, San Diego.

The authors would also like to thank Adriana Di Martino, Michael P Milham and all involved in the collection and aggregation of the ABIDE and ABIDEII datasets.

## Supplemental Materials

**Table S1:** PING demographics by site.

| Site | n | age (range) | Male |
| --- | --- | --- | --- |
| a | 112 | 15.14 (4.08-21.0) | 57 (50.8) |
| b | 105 | 14.69 (3.75-21.0) | 59 (56.2) |
| c | 118 | 12.29 (3.17-21.0) | 58 (49.2) |
| d | 124 | 12.03 (3.17-20.92) | 68 (54.8) |
| e | 64 | 13.42 (4.17-20.92) | 34 (53.1) |
| f | 129 | 8.79 (3.25-20.25) | 65 (50.4) |
| g | 116 | 10.87 (3.42-21.00) | 63 (54.3) |
| **Total** | 768 | 12.28 (3.17-21.0) | 404 (52.6) |

**Table S2:**
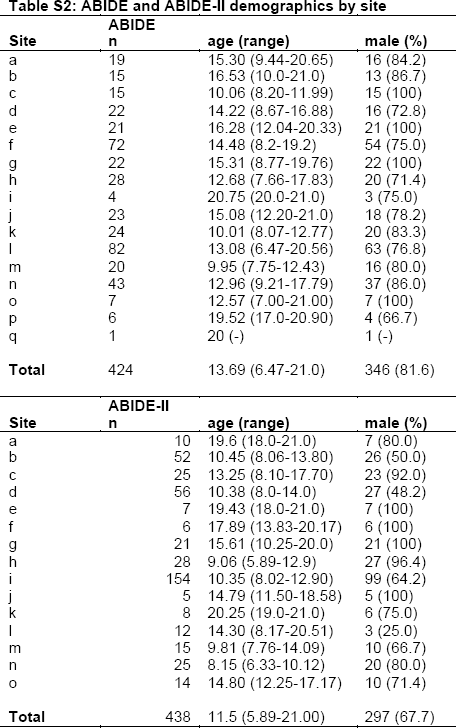
ABIDE and ABIDE-II demographics by site.

| ABIDE |  |  |  |
| --- | --- | --- | --- |
| Site | n | age (range) | male (%) |
| a | 19 | 15.30 (9.44-20.65) | 16 (84.2) |
| b | 15 | 16.53 (10.0-21.0) | 13 (86.7) |
| c | 15 | 10.06 (8.20-11.99) | 15 (100) |
| d | 22 | 14.22 (8.67-16.88) | 16 (72.8) |
| e | 21 | 16.28 (12.04-20.33) | 21 (100) |
| f | 72 | 14.48 (8.2-19.2) | 54 (75.0) |
| g | 22 | 15.31 (8.77-19.76) | 22 (100) |
| h | 28 | 12.68 (7.66-17.83) | 20 (71.4) |
| i | 4 | 20.75 (20.0-21.0) | 3 (75.0) |
| j | 23 | 15.08 (12.20-21.0) | 18 (78.2) |
| k | 24 | 10.01 (8.07-12.77) | 20 (83.3) |
| l | 82 | 13.08 (6.47-20.56) | 63 (76.8) |
| m | 20 | 9.95 (7.75-12.43) | 16 (80.0) |
| n | 43 | 12.96 (9.21-17.79) | 37 (86.0) |
| o | 7 | 12.57 (7.00-21.00) | 7 (100) |
| p | 6 | 19.52 (17.0-20.90) | 4 (66.7) |
| q | 1 | 20 (-) | 1 (-) |
| **Total** | 424 | 13.69 (6.47-21.0) | 346 (81.6) |

| ABIDE-II |  |  |  |
| --- | --- | --- | --- |
| Site | n | age (range) | male (%) |
| a | 10 | 19.6 (18.0-21.0) | 7 (80.0) |
| b | 52 | 10.45 (8.06-13.80) | 26 (50.0) |
| c | 25 | 13.25 (8.10-17.70) | 23 (92.0) |
| d | 56 | 10.38 (8.0-14.0) | 27 (48.2) |
| e | 7 | 19.43 (18.0-21.0) | 7 (100) |
| f | 6 | 17.89 (13.83-20.17) | 6 (100) |
| g | 21 | 15.61 (10.25-20.0) | 21 (100) |
| h | 28 | 9.06 (5.89-12.9) | 27 (96.4) |
| i | 154 | 10.35 (8.02-12.90) | 99 (64.2) |
| j | 5 | 14.79 (11.50-18.58) | 5 (100) |
| k | 8 | 20.25 (19.0-21.0) | 6 (75.0) |
| l | 12 | 14.30 (8.17-20.51) | 3 (25.0) |
| m | 15 | 9.81 (7.76-14.09) | 10 (66.7) |
| n | 25 | 8.15 (6.33-10.12) | 20 (80.0) |
| o | 14 | 14.80 (12.25-17.17) | 10 (71.4) |
| **Total** | 438 | 11.5 (5.89-21.00) | 297 (67.7) |

**Figure S1:**
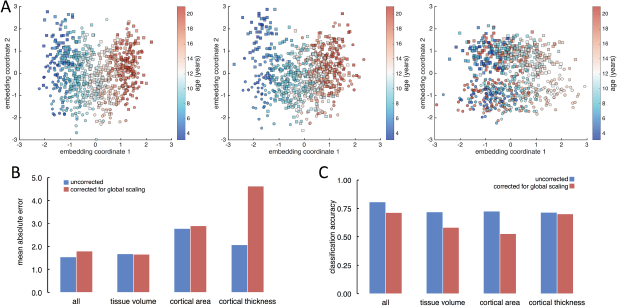
Manifold embedding after correction for global scale. NPE was repeated in the PING dataset after removing variation due to ICV, total surface area and mean cortical thickness from the imaging data. The first two embedding coordinates for each metric are shown in A. Age-related variation remains apparent in both volume and cortical area (left and middle), whereas variation due to sex but not age is preserved in cortical thickness (right). Mean absolute errors for age prediction (B) and classification accuracies for sex prediction (C) are shown below, for the combined embedding and for each metric individually. In general, the combination of all metrics resulted in an improved performance over single metrics.

**Figure S2:**
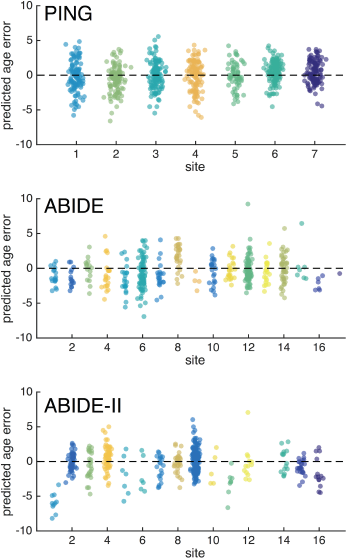
Site variation in age prediction error for PING, ABIDE and ABIDE-II. Predicted age error is shown within individual sites for each cohort.

**Figure S3:**
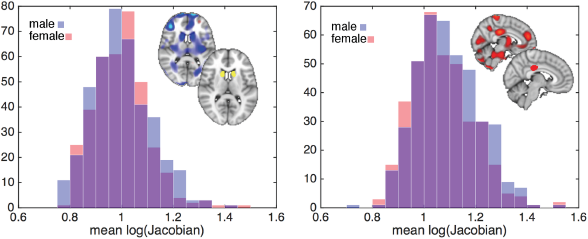
Regional sex differences show a small effect size. Two regions are highlighted, based on the coefficient image in Fig 2A, where larger volume is associated with female sex (left: negative model coefficients, caudate mask inset) and male sex (right: positive model coefficients, posterior cingulate mask inset). Histograms show the volume distribution (measured as the mean log Jacobian within each mask) across sexes. In a univariate setting, the average difference in volume in these regions is small. This highlights how appropriate multivariate subspace projection methods are able to robustly isolate structured variation within populations, even if the effect size within individual regions is small.

